# Computationally profiling peptide:MHC recognition by T-cell receptors and T-cell receptor-mimetic antibodies

**DOI:** 10.1101/2022.07.15.500208

**Authors:** Matthew I. J. Raybould, Daniel A. Nissley, Sandeep Kumar, Charlotte M. Deane

## Abstract

T-cell receptor-mimetic antibodies (TCRms) targeting disease-associated peptides presented by Major Histocompatibility Complexes (pMHCs) are set to become a major new drug modality. However, we lack a general understanding of how TCRms engage pMHC targets, which is crucial for predicting their specificity and safety. Several new structures of TCRm:pMHC complexes have become available in the past year, providing sufficient initial data for a holistic analysis of TCRms as a class of pMHC binding agents. Here, we profile the complete set of TCRm:pMHC complexes against representative TCR:pMHC complexes to quantify the TCR-likeness of their pMHC engagement. We find that intrinsic molecular differences between antibodies and TCRs lead to fundamentally different roles for their heavy/light chains and Complementarity-Determining Region loops during antigen recognition. The idiotypic properties of antibodies may increase the likelihood of TCRms engaging pMHCs with less peptide selectivity than TCRs. However, the pMHC recognition features of some TCRms, including the two TCRms currently in clinical trials, can be remarkably TCR-like. The insights gained from this study will aid in the rational design and optimisation of next-generation TCRms.

## Introduction

The human adaptive immune system relies upon B-cells and T-cells that use characteristic membrane-bound immunoglobulins, B-cell receptors (BCRs) and T-cell receptors (TCRs), to recognise a broad range of pathogenic antigens, many of which are proteinaceous. BCRs, and their secreted soluble analogues, antibodies, recognise complete soluble or membrane-bound extracellular proteins. T-cell receptors (TCRs), meanwhile, are focused through thymic development to recognise fragments of intracellularly- or extracellularly-derived peptides presented on cell surfaces by either a class I or class II polymorphic major histocompatibility complex (pMHCs) (1).

Despite their different natural roles, the binding domains of antibodies and TCRs bear several commonalities. They are both comprised of two analogously gene-recombined chains (termed ‘heavy/light’ (H/L) and ‘beta/alpha’ (B/A) for antibodies and TCRs respectively) and six complementarity-determining region (CDR) loops that together constitute most of their binding sites. These similarities have long motivated efforts to understand whether antibodies can engage pMHCs with TCR-like specificity (2). ‘TCR-mimetic antibodies’ (TCRms) that specifically recognise fragments of the intracellular proteome could offer pinpoint recognition of aberrant cells, transforming immunohistochemistry and immunotherapy.

TCRms also offer a number of practical advantages over TCRs in terms of soluble drug development. TCR:pMHC binding affinities lie in the 1-100*μ*M range (3, 4), meaning they must be affinity-engineered for use as a monovalent binding arm that recognises low copy number pMHCs (5). By contrast, antibody:antigen monovalent binding frequently occurs at the required range of affinities for therapeutic effect (low nM-pM) (6, 7). Therapeutic antibody development pipelines are also more established than their TCR equivalents, facilitating TCRm clinical translation and adaptation to multispecific formats exploiting proven cancer immune-modulation and T-cell redirection strategies (8–11).

Early studies sought to elicit natural TCRms *via* allogenic mouse immunisation and established that BCRs can be raised against non-self peptide:non-self MHC complexes. Though most antibodies were able to engage the MHC regardless of presented peptide, a smaller fraction were peptide-dependent (*i.e*. at least somewhat TCR-mimetic) (12–15).

TCRm isolation strategies shifted towards the use of *in vitro* phage-display libraries (14, 16), both sidestepping the need to account for self-tolerance and enabling rounds of positive and negative selection to enrich for stronger, more peptide-dependent binding. By 2020, these libraries had produced a variety of TCRms against a wide range of both class I and class II pMHC targets (15, 17–20), although the extent of their peptide specificity, and thus the breadth of their applicability, was still highly variable. Most have been used as chemical probes, for which the required peptide specificity is lower than that required of a therapeutic administered across heterogeneous (MHC-compatible) populations.

Experimental peptidome binding assays performed on three early-generation TCRms (17, 19, 20) suggested that they were unlikely to achieve the levels of specificity required for therapeutic applications (21, 22). However, several TCRms with apparently high specificity have been reported in the past year, fostering renewed interest in this therapeutic modality and resulting in a more than doubling of the number of crystal structures of TCRm:pMHC complexes (23–28). These include an anti-Wilms’ Tumor Antigen 1 (WT1) TCRm (23) and an anti-alpha ferroprotein (AFP) TCRm (28) that have both progressed to clinical trials (23). Two neoantigen peptide:MHC-specific TCRms were also identified that achieve complete selectivity over their wildtype peptide equivalents each differing by just a single residue mutation (24, 25).

Here, we harness this recent increase in structural data on TCRm:pMHC complexes to computationally dissect their molecular recognition properties. We outline where and why TCRm:pMHC binding features tend to align with and differ from representative antibody:antigen and TCR:pMHC complexes. High-throughput interaction profiling of static complexes reveals that molecular differences between antibodies and TCRs result in differential CDR involvement in the pMHC binding event. This tends to lead to more variable peptide sensitivity, but does not preclude some TCRms from recognising pMHCs with similar features to those seen across TCRs. We also perform all-atom simulations which reveal that energetic hotspots in the MHC can play a key role in TCRm binding. TCRs seem to avoid this behavior, instead reliably exploiting energy hotspots on the peptide surface. Finally, we highlight TCR-like pMHC recognition features in the first TCRms to achieve sufficient specificity to reach the clinic (11D06 (23) and AFP-TCRm (28)) *versus* a TCRm with several known off-targets (ESK1). Overall, our analysis begins to quantify TCR-likeness across TCRms, enabling rational TCRm selection, optimisation, and design based on the natural cognate partners of pMHCs.

## Results

We began our analysis by identifying sets of representative antibody:antigen, TCR:pMHC and TCRm:pMHC complexes from SAbDab (6, 7) and STCRDab (4) (see Methods).

We found twelve pMHC-binding antibodies, eleven of which have binding modes that transect the peptide binding groove (17–20, 23–26) and one alloantibody (29) that contacts only the MHC and so was not classified as a TCRm. After filtering for interface redundancy (see Methods), we identified 10 representative TCRm:pMHC complexes (9 MHC class I, 1 MHC class II), 60 representative TCR:pMHC complexes (52 MHC class I, 8 MHC class II), and 824 representative antibody:antigen complexes. Unless otherwise stated, properties of TCRs or TCRms engaging MHC class I and class II are pooled as they are both engaged by the same genetic class of TCR (*αβ*).

### pMHC complexes offer an unusually broad binding surface that is engaged differently by TCR and TCRm CDRs

To quantitatively compare the properties of immunoglobulin:antigen interfaces, we computed the buried surface area (BSA) and patterns of formal interactions across our representative complexes (see Methods, SI Methods).

#### Global Interface Properties

Calculating BSA over the whole immunoglobulin:antigen interface (Fig. 1A) reveals that pMHC binding events result in atypically broad interfaces relative to general antigen complexes (TCRs *μ*: 1852.6Å^2^, sd: 243.8Å^2^ and TCRms *μ*: 2015.7Å^2^, sd: 183.9Å^2^; *versus* general antibodies *μ*: 1496.7Å^2^, sd: 468.7Å^2^).

**Fig. 1.**
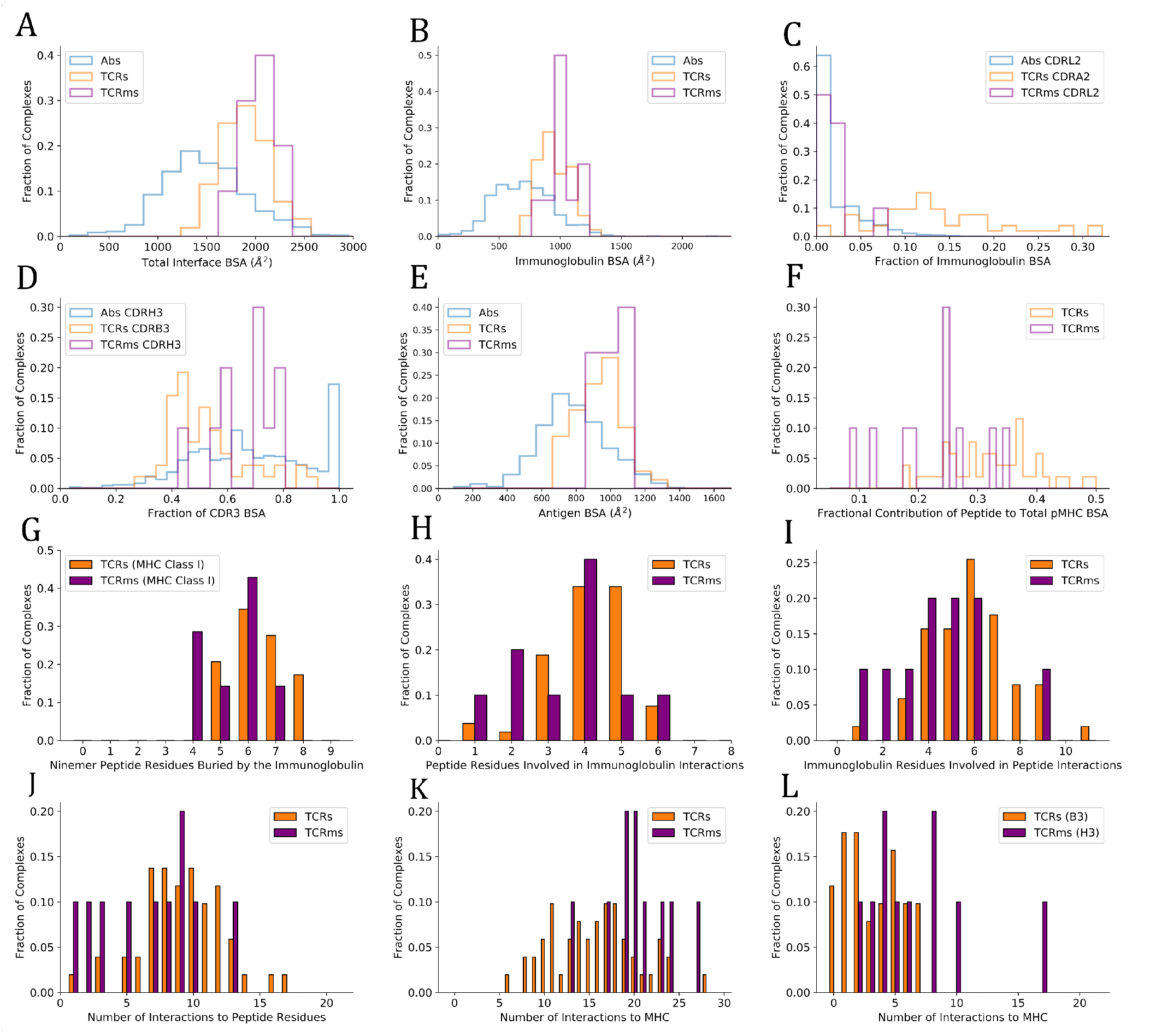
Buried surface area (BSA) and interaction profiles across the representative complexes from the Structural Antibody Database (6, 7) and Structural T-cell Receptor Database (4). A: The total BSA across the immunoglobulin:antigen interface. B: The immunoglobulin portion of the total interface BSA. C: The fractional contribution of CDRL2 (antibodies, TCRms) or CDRA2 (TCRs) to immunoglobulin BSA. D: The fractional contribution of CDRH3 (antibodies, TCRms) or CDRB3 (TCRs) to CDR3 BSA. E: The antigen/pMHC portion of the total interface BSA. F: The fractional contribution of peptide BSA to pMHC BSA. G: The number of peptide residues buried in each immunoglobulin to ninemer peptide:MHC Class I complexes. H: The number of peptide residues involved in binding interactions to the TCRms/TCRs. I: The number of immunoglobulin residues involved in binding interactions to the peptide across the TCRms/TCRs. J: The number of interactions between the immunoglobulin and the peptide across the TCRms/TCRs. K: The number of interactions to MHC residues across the TCRms/TCRs. L: The number of interactions to MHC residues made by the CDRH3 (TCRms) or CDRB3 (TCRs) loop.

The flat pMHC topology appears to place constraints on the CDRH3/CDRB3 length of their cognate immunoglobulins. Though assembled *via* a common VDJ recombination mechanism, TCR CDRB3s only have a length range of 10-17 (30), while antibody CDRH3s span lengths of 5-30+ (31–33). The TCRm CDRH3s range from length 10 to length 16, biased to the lower end of the range sampled in natural antibodies, with a mean value closer to that seen across TCRs, and entirely within the relatively narrow band of TCR CDRB3 or CDRA3 loop lengths (30, 34) (SI Table 2). This suggests that shorter CDR3 lengths render TCRs/TCRms unable to achieve sufficient interactions to bind a pMHC, while longer lengths may result in destabilising clashes with the pMHC surface.

A residue-level interaction analysis of the immunoglobulin:pMHC complexes shows that the broad interface comprises different total numbers of interactions in TCRs and TCRms (*μ*: 23.2, sd: 6.2; *μ*: 27.0, sd: 3.4 respectively), however both classes of immunoglobulin use a similar balance of hydrophobic, aromatic, and polar interactions (SI Table 1).

#### Binding Properties by Immunoglobulin Region

Only considering the immunoglobulin contribution to the BSA, the TCRm profile (*μ*: 1015.8Å^2^, sd: 111.0Å^2^) is again closer to typical TCRs (*μ*: 934.4Å^2^, sd: 127.2Å^2^) than typical antibodies (*μ*: 707.5Å^2^, sd: 260.0Å^2^) (Fig. 1B).

However, dissecting this BSA by CDR contributions demonstrates that antibody and TCR CDR loops play different roles in antigen binding. For example, antibody CDRL2 loops lie unburied in over 50% of general antigen complexes while TCR CDRA2 loops are buried in the pMHC interface to a much greater extent, a greater proportion of the time (Fig. 1C).

Equally, when considering the relative contributions of CDR3s to pMHC recognition, we find that TCRm binding tends to be biased towards burial of CDRH3 and away from CDRL3, more typical of general antibody:antigen complexes (Fig. 1D), while TCRs exploit their CDR3 loops more evenly, if anything with a slight bias towards the CDRA3 loop (the genetic equivalent to CDRL3).

The differences in CDR usage in pMHC binding can be related to the fact that some CDR loops have markedly different length preferences in antibodies than TCRs. For example, antibody CDRH2s and CDRL2s have median IMGT lengths of 8 and 3, respectively (31). In contrast, TCR CDRB2 and CDRA2 loops have median lengths of 6 and 5 (34). Similarly, while antibody CDRH3 and CDRL3 loops have a median length of 15 and 9, TCR CDRB3 and CDRA3 loops have median lengths of 12 and 13 (30, 31, 34). The more even balance in CDR lengths between equivalent CDR loops on the VA and VB chains is consistent with the lower observed bias towards VDJ-chain dominated binding (Fig. 1D).

In summary, the differences in the molecular properties of antibodies and TCRs have a direct impact on the roles of their CDR loops during pMHC recognition, with no apparent functional link between antibody/TCR chains made by analogous gene recombination mechanisms.

#### Binding Properties by Antigen Region

BSAs computed only across the antigen (Fig. 1E) reveal that the expected result that TCRs (*μ*: 918.2Å^2^, sd: 122.0Å^2^) and TCRms (*μ*: 999.8Å^2^, sd: 79.5Å^2^) tend to bury a larger total area of the antigen interface relative to general antibodies (*μ*: 789.0Å^2^, sd: 222.7Å^2^).

Although TCRs and TCRms bury a similar area of the pMHC, splitting this area into contributions by the peptide and MHC reveals that the TCRms are strongly biased towards a larger MHC BSA, with the effect that TCRms tend to recognise a smaller proportion of peptide surface during their pMHC recognition events (Fig. 1F). This can also be expressed as the total number of peptide residues buried at least to some extent. Regardless of peptide length, no cognate TCR or TCRm has yet been found that can bury every peptide residue. However, for example in ninemer peptides presented by MHC class I, burial of eight peptide residues is not uncommon across TCRs (17.2%) but has not yet been observed in any TCRms (Fig. 1G). Only 1/7 (14.3%) ninemer-pMHC class I-binding TCRms buries seven peptide residues (11D06 [7BBG]), while at least this many peptide residues are buried in 13/29 (44.8%) of corresponding TCRs.

Though we observe that TCRms have a lower proportion of pMHC BSA from the peptide, they do engage a similar number of peptide residues using a similar number of immunoglobulin residues as TCRs (Figs. 1H, 1I). This also results in a similar number of formal interactions (Fig. 1J). However, the BSA signal that TCRm:pMHC recognition is disproportionately biased towards MHC is recapitulated in the fact that TCRms tend to make more formal interactions to MHC residues than do TCRs (Fig. 1J). Dissecting these contributions by immunoglobulin region shows this predominantly originates in TCRm CDRH3s being less peptide-focused than TCR CDRB3 loops (Fig. 1K). These profiles suggest that, on aggregate, the current set of isolated TCRms are likely to be less peptide-selective than typical *in vitro/in vivo*-selected TCRs.

### TCRms can approach pMHCs with a diagonal orientation, but this does not guarantee TCR-like pMHC recognition

It has been previously shown that TCRs converge around diagonal engagement of pMHCs with the centre of mass of the beta chain sitting over the C-terminus of the MHC *α*_1_ helix and the centre of mass of the alpha chain sitting over the C-terminus of the MHC *α*_2_ helix (15, 35). This is quantified by the ‘docking/crossing angle’, calculated as the intercept of the line connecting conserved centres of mass within the variable region and the axis of the peptide binding groove. Our 60 representative TCR:pMHC complexes almost entirely comply with this canonical binding definition (*μ*: 45.8°, sd: 17.0°, SI Fig. 2A), with one outlier (Protein Data Bank identifier (PDB ID) 4Y19 (36)) (37) that engages pMHC with a diagonal but reverse polarity mode (136.7°; *i.e*. the VB chain sits above the MHC *α*_2_ helix while the VA chain occludes the MHC *α*_1_ helix, SI Fig. 2B).

To visualise their binding orientations, we aligned our set of TCRm complexes along the canonical peptide binding groove axis (Fig. 2). While two of the TCRms bound in a non-diagonal fashion, the rest (80% of non-redundant TCRms) adopted a diagonal pMHC binding mode that fell within the range of absolute docking angles set by TCRs (Fig. 2). However, in contrast to the TCRs, TCRm diagonal pMHC binding is more frequently achieved using reverse polarity (where VH is in the position of VA and VL is in the position of VB), reinforcing the notion that antibody and TCR chains with similar gene recombination mechanisms are not necessarily analogous in terms of pMHC recognition.

**Fig. 2.**
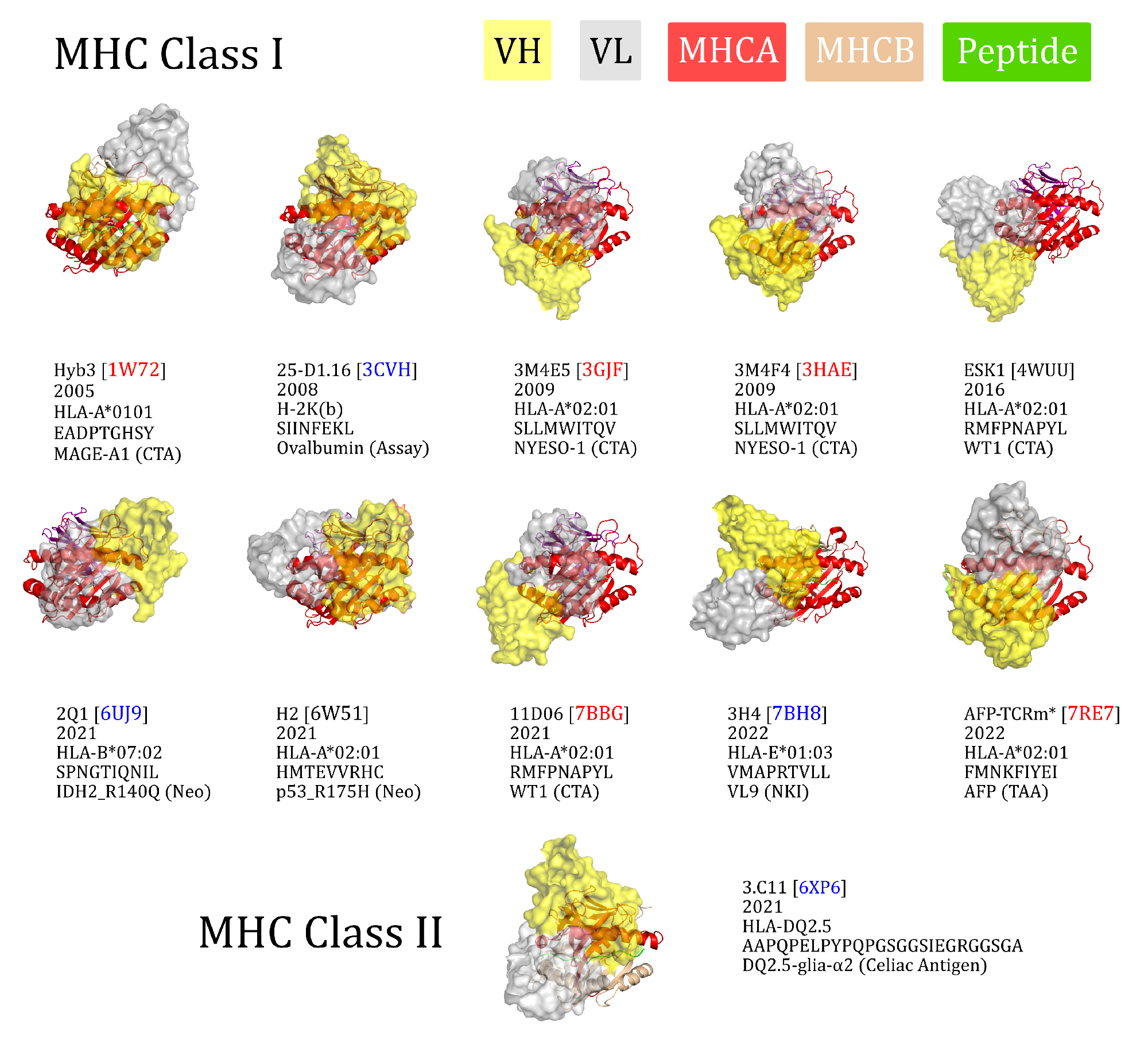
The eleven TCR-mimetic antibodies (TCRms) with solved structures in complex with their cognate pMHC, aligned by the MHC residues. The perspective set so that the x-axis is the vector through the C_*α*_ atoms of the peptide anchor residues. 3M4E5 and 3M4F4 are closely related and derive from the same screening campaign, so are jointly represented by 3M4E5 in the analysis. Metadata is supplied below each structure: the TCRm name and Protein Data Bank (PDB) identifier, year of release in the PDB, MHC allele, antigen amino acid sequence, and antigen common name. ESK1 and H2 fall outside the range of diagonality seen in TCR:pMHC binding (PDB codes written in black). Diagonally-binding TCRms with VH in the position of VB and VL in the position of VA (canonical polarity binders) have blue PDB codes, while those with VH in the position of VA and VL in the position of VB (reverse polarity binders) have red PDB codes. CTA: cancer testis antigen, MHCA: MHC alpha chain, MHCB: MHC beta chain, Neo: neoantigen. *We have labelled the TCRm from 7RE7 as AFP-TCRm as it has no name in the seminal paper (28).

Convergence upon diagonal pMHC binding across TCRs is thought to be driven by improved typical TCR specificity, achieved by positioning the most hypervariable loops within interaction distance of the peptide, the key locus of pMHC variability (34). We therefore surveyed the properties of diagonally *versus* non-diagonally engaging TCRms to investigate to what extent this property correlates with more TCR-like pMHC recognition (SI Fig. 1).

The property distributions indicate that TCRm diagonal engagement is not systematically linked with total pMHC BSA (SI Fig. 1E) nor a higher fraction of peptide contact area (SI Fig. 1F). Some diagonal modes result in few formal interactions between the TCRm CDR3s and the peptide (e.g. Hyb3, with just two interactions to the peptide from CDRH3), while others result in numbers large even by TCR standards (Fig. 1J; 3M4E5 has three formal interactions to the peptide from CDRH3 and seven from CDRL3). However, one property that appears to be systematically linked to TCRm diagonal engagement is the ability to bury a greater number of peptide residues; where N is the total number of peptide residues, non-diagonal pMHC class-I modes contact a maximum of N-4 residues, while diagonal modes frequently bury more, up to a currently-observed maximum of N-2 (SI Fig. 1G).

Overall these mixed results show that not all diagonal TCRm pMHC binding modes yield TCR-like pMHC engagement profiles, and that this property alone is likely insufficient for capturing the specificity of a TCRm.

### TCRms and TCRs have different trends in binding energetics even in identical pMHC contexts

We next performed molecular dynamics studies on three pMHC contexts for which we have a crystal structure in the PDB of at least one TCRm partner and at least one natural/affinity-enhanced TCR partner, allowing us to investigate their binding energetics. We selected three cases studies in which HLA-A*0201 presents a different peptide antigen:

1. Wilms’ Tumor 1 (WT1) antigen (RMFPNAPYL), bound to two TCRms and one affinity-enhanced TCR; ESK1 (PDB ID: 4WUU), 11D06 (PDB ID: 7BBG), and a7b2 (PDB ID: 6RSY), respectively.
2. New York esophageal squamous cell carcinoma 1 (NY-ESO-1) antigen (SLLMWITQV), bound to one TCRm, one natural TCR, and three affinity-enhanced TCRs; 3M4E5 (PDB ID: 3GJF), sp3.4 (PDB ID: 6Q3S), NYE_S1 (PDB ID: 6RPB), NYE_S2 (PDB ID: 6RPA), and NYE_S3 (PDB ID: 6RP9), respectively.
3. p53_R175H neoantigen (HMTEVVRHC), bound to one TCRm and three natural TCRs; H2 (PDB ID: 6W51), 1a2 (PDB ID: 6VQO), 12-6 (PDB ID: 6VRM), and 38-10 (PDB ID: 6VRN), respectively.

To characterize the energetic differences between the bound and unbound states of each complex, we ran sets of thirty 5-ns all-atom explicit solvent molecular dynamics simulations, using Molecular Mechanic Generalized-Born Surface Area (MMGBSA) (39–41) to compute overall free-energy changes upon binding as well as the per-residue contributions from both the pMHC and TCR/TCRm (see Methods, SI Methods, SI Figs. 3-5). We used an ensemble sampling approach, *i.e*. a set of short statistically independent simulations initiated with different random velocities from the same starting structure, to ensure thorough sampling of the bound configurations.

Quantifying the relative contributions of the peptide, MHC *α*1 helix, and MHC *α*2 helix to the overall free energy change of the pMHC (see SI Methods for equations), we find that both natural TCRs and affinity-enhanced TCRs tend to gain a larger proportion of their binding energy through the peptide than do TCRms (Fig. 3, affinity-enhanced TCRs labelled TCR*). Within a given pMHC case study, no single TCRm achieved a higher proportion of binding energy from the peptide than any TCR. It is striking that some peptide antigens appear more tractable than others in terms of achieving a significant contribution to the overall binding energy. For example, even an affinity-enhanced TCR engaging the WT1 pMHC only reaches around 40% peptide contribution to binding free energy, while natural tumor-infiltrating lymphocytes complementary to p53_R175H achieve up to 65%. We also considered the energies contributed by individual residues to identify interaction hotspots (Table 1, SI Figs. 2-4, SI Table 3). Arbitrarily, we define interaction hotspots as individual residues that have a predicted free energy of -7 kcal/mol or stronger. All the natural TCRs assessed have at least one hotspot residue in the peptide, suggesting this may be a frequent feature of natural peptide recognition and consistent with current hypotheses on TCR recognition (42, 43). That the three p53_R175H specific TCRs found in tumorinfiltrating lymphocytes achieve more neoantigen recognition at R7 rather than H8 (the site of somatic mutation) shows that this binding hotspot does not necessarily lie at the residue that distinguishes self from non-self.

**Fig. 3.**
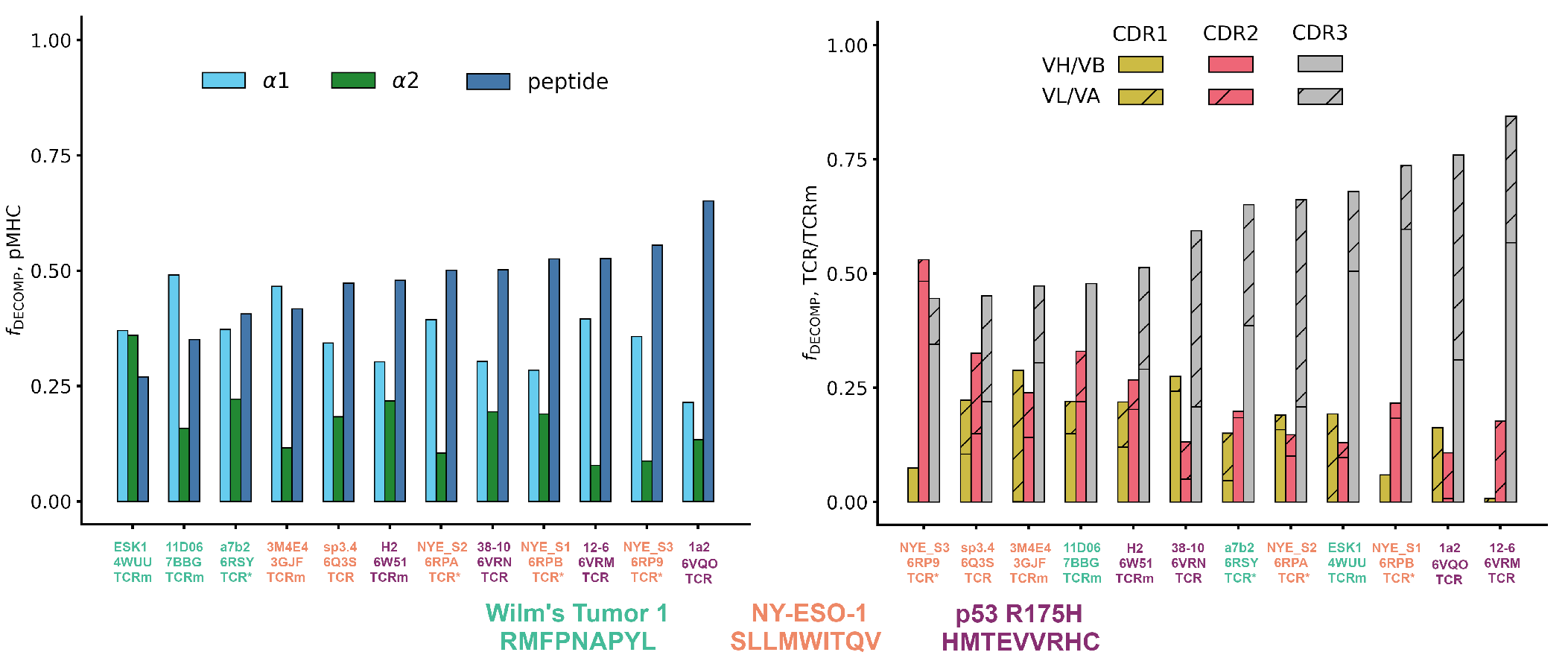
The energetics of pMHC binding of TCR-mimetic antibodies (TCRms), natural T-cell receptors (TCRs) and affinity-enhanced T-cell Receptors (TCR*s). (LHS) Relative contributions of the peptide, the MHC *α*1 helix, and the MHC *α*2 helix to the MMGBSA free energy change (38). Individual values were computed with SI Equations 1-3, and then rank ordered by increasing peptide contribution. (RHS) Relative contributions to binding free energy of the CDR1, CDR2, and CDR3 loops for the heavy/beta chain (no hatch) and light/alpha chain (diagonal hatch), calculated as described in Methods. Complexes were rank ordered by increasing CDR3 contribution. Note that the CDR1, CDR2, and CDR3 bars do not sum to exactly one for all complexes as small negative values of the fraction of the DECOMP energy (*f_decomp_*), representing CDRs that experience repulsive interactions with the pMHC, were passed to zero for visual clarity. NY-ESO-1: New York esophageal squamous cell carcinoma 1.

**Table 1.**
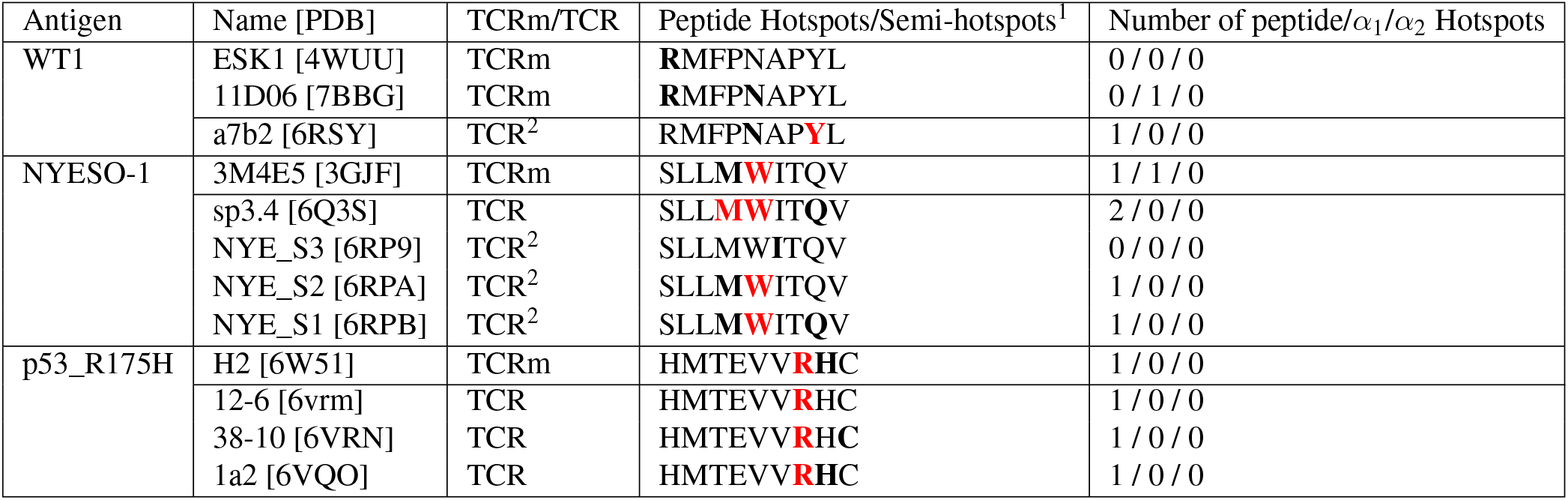
Decomposed per-residue energetic profiles for the case study TCR and TCRm complexes, grouped by pMHC region (peptide, MHC *α*_1_ helix, or MHC *α*_2_ helix). ^1^‘Hotspots’ are defined as residues predicted to have an attractive per-residue contribution of ≥ 7 kcal/mol to free energy based on DECOMP analysis (bold red text). ‘Semi-hotspots’ are defined as residues that contribute between -4 and -7 kcal/mol (bold black text). ^2^Affinity-enhanced TCRs.

By contrast, only TCRms were found to have hotspots that lie within the MHC (3M4E5 and 11D06). Despite their origins in different light chain loci (IGLV2-11 and IGKV1-5, respectively), the hotspot occurs at the same residue in the *α*1 helix (R65), which in both cases forms a salt bridge to D56 of CDRL2. That such non-peptide hotspots were not readily observed in TCR formats provides further evidence that current TCRms may be prone to higher inherent affinity for the MHC and thus poorer typical peptide specificity.

Finally, to characterize the differences in the energetics of CDR binding to the pMHCs, we computed the fractional contributions of each CDR to the total CDR free energy change (Fig. 3). All TCRms and most TCR contexts were CDR3-dominated in their binding energetics. The balance of CDR[H/B]3 to CDR[L/A]3 energy reflected the picture seen in the general analysis of interface properties: TCRms gain more binding energy through CDRH3 than CDRL3, while in TCRs either CDR3 can be energetically dominant.

## Discussion

TCRs exhibit a range of pMHC specificities, with a degree of polyspecificity considered an advantageous property to maximise the TCR repertoire’s breadth of antigen recognition (44). However, whether due to evolutionary constraints or thymic selection mechanisms, TCRs have been observed to converge around several pMHC recognition properties including a canonical docking polarity, orientation, and the positioning of their most variable CDR loops above the peptide binding groove of the MHC (45). Regardless of underpinning mechanisms, this suggests there are conserved properties of pMHC recognition that grant TCRs the basal level of peptide specificity necessary to avoid widespread autoimmunity *in vivo*. The properties of their interfaces should therefore shed light on pMHC recognition features that ought to be generally advantageous to other molecules seeking to mimic them. In this paper we have shown that, despite their broader genetic similarities, the idiotypic molecular configurations of antibody and TCR CDR loops contribute to the observed differences in their pMHC recognition tendencies. So far we lack evidence that genetically-equivalent antibody CDR loops can precisely imitate the contact profiles of TCR CDR loops in their canonical binding footprint. However, relaxing the requirement for loop and chain equivalency, TCRms do exhibit a spectrum of TCR-likeness in the way they engage pMHC.

Within this spectrum, it is currently difficult to set thresholds for how TCR-like a TCRm needs to be to be clinically viable. Three of the TCRms, ESK1, 11D06, and AFP-TCRm, may begin to shed light on the answer. ESK1 is potentially the least TCR-like TCRm analysed in this study, engaging the pMHC with an orthogonal binding mode (Fig. 2), the lowest fraction of binding energy to the peptide of any simulated complex (Fig. 3), and several burial/interaction profiles on or outside of the bounds seen previously in TCRs (SI Fig. 6). By contrast, 11D06 and AFP-TCRm are considerably more TCR-like, engaging the pMHC diagonally (Fig. 2) and with far fewer MHC interactions than ESK1 (SI Fig. 6). 11D06 buries the highest number of peptide residues, and AFP-TCRm the highest fraction of peptide/pMHC surface area, of any class-I TCRm to date.

Though ESK1 was reported many years before 11D06 and AFP-TCRm, it has not yet progressed through preclinical development and has several known human proteome off-targets (22), while 11D06 and AFP-TCRm have already advanced to in-human clinical trials. This suggests that increased TCR-likeness may be beneficial for clinical progression. However, it is worth reiterating that even these clinical TCRms are not perfectly TCR-like. For example, 11D06 harbors an interaction hotspot in the MHC, has a smaller percentage of pMHC binding energy from the peptide than any TCR assessed in the simulation study, and its VJ-recombined CDR3 loop plays no role in binding. This implies that some TCR-like pMHC recognition properties could be more important than others for therapeutic development.

One repeating feature of the three TCRms able to recognise a large proportion of peptide residues without extensive MHCA interaction (3M4E5 and both clinical-stage TCRms) is a salt bridge between IMGT residue D56 at the start of CDRL2 and residue R65 on the MHCA *α1* helix. While this salt bridge seems to result in an energetic hotspot on the MHC (SI Figs. 3-4), this may be compensated for by helping to orient the TCRm in a highly peptide-sensitive ‘reverse polarity’ pMHC recognition mode. It may therefore be worth further investigation as to whether this recurrent interaction can be exploited in future TCRm drug development pipelines. There are several limitations to this study, primary amongst which is the relative paucity of TCRm structures, and particularly pMHC contexts, upon which to make conclusions about general TCRm pMHC recognition. Selection bias that cannot be accounted for through non-redundancy filtering alone is also likely to influence our conclusions. For example, it is likely that TCRms that have more promising specificity profiles would be more likely to be subjected to crystallographic analysis, which may artificially increase their apparent average TCR-likeness. The TCR-likeness of TCRms may also be expected to increase over time as *in vitro* negative selection techniques become ever more rigorous (e.g. negative baiting against panels of molecular mimicking off-targets, single point mutants of the target, or general broad screening panels of representative self or artificial peptides loaded onto MHC). Additionally, a notable trend in recent years has been to solve TCR:pMHC complexes that ‘break the rules’ rather than adhere to them (46), which may interfere with interpretability of the properties of our ‘representative TCRs’ as reflective of typical TCR behavior.

Nevertheless even with these early data, some general properties of TCRms and TCRs, such as their balance of peptide *versus* MHC recognition, show marked differences. As more TCRms and immune mobilising monoclonal T-cell receptors against cancer (immTACs) are developed and structurally characterized, it should be insightful to evaluate their binding properties and assess which pMHC recognition features are most reliably linked with successful clinical progression. For now, our TCR:pMHC complex profiles offer an initial set of benchmarks for the rational computational derisking of novel TCR-mimetic modalities.

## Methods

### Numbering and Region Definitions

To enable a direct comparison between antibodies and TCRs, and between heavy and light chains, the IMGT numbering scheme and CDR definitions were used throughout this work (CDR1: IMGT residues 27-38, CDR2: IMGT residues 56-65, CDR3: IMGT residues 105-117 (47)). ANARCI was used to number all sequence inputs (48). Where necessary, MHC chains were renumbered to enable a direct comparison between TCRm and TCR complexes.

### Structure Datasets

SAbDab (4, 6) and STCRDab (4) were downloaded on 30^th^ September, 2022. All complexes were stripped of explicit hydrogens, heteroatoms, and water molecules, and only immunoglobulins with protein antigens were considered. These databases were then mined for particular subcategories as follows:

1. TCR-mimetic antibody (TCRm) complexes were identified by filtering SAbDab with the search terms ‘HLA’ and ‘MHC’, followed by manual validation.
2. Representative sets of non-redundant high quality antibody:antigen and TCR:pMHC (*αβ* only) complexes were derived by first filtering SAbDab and STCRDab for structures of complexes solved by X-ray crystallography to ≤ 2.5 Å resolution. For each class of immunoglobulin separately, the IMGT-defined CDR sequences (47) were concatenated in a consistent order and used as inputs to greedy clustering by cd-hit (49) (80% sequence identity threshold), to create ‘paratope clusters’. To identify cases where chemically-similar paratopes bind to significantly different antigens, we performed a second round of clustering over the paratope clusters using the concatenated sequence of the antigen(s) associated with each antibody (concatenated in descending length order), or the sequence of the presented peptide for the TCRs. cd-hit was run at an 80% sequence identity threshold, allowing a minimum alignment length of as little as 20%. This ensured truncated antigens were not considered as different targets (e.g. an anti-coronavirus antibody solved in complex with the receptor-binding domain would be considered the same context as a complex of the same antibody binding to the full-length spike protein). This resulted in sets of 824, 52, and 8 representative antibody:antigen, TCR:pMHC (class 1) and TCR:pMHC (class 2) complexes respectively.
3. Complexes of TCRs and TCRms engaging the same peptide were obtained by searching SAbDab and STCRDab by antigen sequence for peptide fragments with 100% sequence identity match.

### Buried Surface Area Calculations

Complete immunoglobulins were separated from their antigens (*i.e* the co-ordinates of the partner were deleted in a copy of the original file), and the difference in solvent-accessible surface area between each residue in the original complex and artificially-generated ‘apo’ state was recorded. Per-residue solvent-accessible surface area was calculated using an inhouse implementation of the Shrake and Rupley algorithm (50), applying a probe radius of 1.4Å.

### Interaction Mapping

The Arpeggio (51) software package was used to assess complexes for hydrophobic, aromatic, hydrogen bond, and salt bridge interactions. A set of rules were established to interpret the arpeggio outputs at a per-residue level (see SI Methods).

### Molecular dynamics simulations and analysis

pMHC-TCR and pMHC-TCRm complexes were prepared in Amber-Tools21 (52) and all simulations run in OpenMM v7.5 (53). Cis peptide bonds and chiral centers with incorrect stereochemistry within rebuilt sections of protein structures were identified and corrected using the Cispeptide and Chirality (54) plugins of Visual Molecular Dynamics (55) v1.8.3. Corrected peptide bonds were maintained in the trans configuration with periodic torsion restraints with force constant 500 kJ/mol. These restraints were removed during the final unrestrained equilibration along with positional restraints on Cα atoms. Complexes were then solvated in orthorhombic boxes with a minimum of 1.4 nm between protein atoms and the box edge. Sodium and chloride ions were added to neutralize each system and bring the salt concentration to 0.15 M.

All simulations were carried out using FF14SB (56) and TIP3P (57) forcefield parameters with a Langevin thermostat (friction coefficient 1 ps^−1^) and, for constant-pressure simulations, a Monte Carlo barostat (pressure 1 bar). Complexes were minimized and then heated to 298 K at constant volume with protein atoms restrained. These restraints were relaxed over a set of five 100-ps simulations at constant pressure, culminating with 100-ps of unrestrained equilibration. Finally, 5-ns production simulations at constant pressure were run. Thirty independent replicas of this protocol were executed for each complex (58). MMGBSA calculations were performed with MMPBSA.py (59) using 100 frames collected every 40 ps from the final 4 ns of each 5-ns trajectory (i.e., 3000 frames per complex). Per-residue energy contributions to the binding energy were computed using the MMPBSA.py DECOMP functionality (59). Further detail on all simulation methods can be found in the SI.

### Visualizations

All manuscript visualizations were created using open-source PyMOL or matplotlib version 3.5.2.

## Data Availability

Datasets are publicly available on Zenodo at 10.5281/zenodo.7220531. The cleaned, filtered, and IMGT-numbered complexes used for the high-throughput immunoglobulin:antigen complex analysis are supplied as Dataset S1. PQR-format structures generated with H++ (59) for simulations of case study pMHC-TCR and pMHC-TCRm complexes are available as Dataset S2.

## Acknowledgements

MIJR and DAN gratefully acknowledge funding through a postdoctoral research grant sponsored by Boehringer Ingelheim. The authors thank Drs. Nicolas Sabarth, Niksa Kastrapeli, Andrew Nixon, Justin Scheer, Noah Pefaur, Srinath Kasturirangan, Paul Adams, Renate Konopitzkey and several other researchers at Boehringer Ingelheim for numerous insightful discussions and their critical reading of this manuscript.

