## Supplementary Information for "Computationally profiling peptide:MHC recognition by T-cell receptors and T-cell receptor-mimetic antibodies"

### Per-residue interaction Analysis with Arpeggio

Arpeggio featurises input protein complexes with predicted interaction types based on the relative distances and orientations of PDB atom types. In our analysis, we only considered the four most attractive interaction types: hydrophobic, aromatic, hydrogen bond (not weak), and salt bridge interactions. To interpret these arpeggio interaction outputs on a per-residue level requires a consolidation of these atom-level interaction profiles, which we performed according to the following post-processing steps:

1. Where two residues had multiple hydrophobic atom-atom contacts this was recorded as one residue-residue "hydrophobic" interaction.
2. Where two residues had multiple aromatic atom-atom contacts this was recorded as one residue-residue "aromatic" interaction.
3. A residue-residue pair can be labelled as contributing both a hydrophobic and an aromatic interaction due to their different electrostatic origins.
4. Polar interactions between oppositely charged residues were initially labelled as 'salt bridge/hydrogen bond'. Once all interactions were recorded the salt bridge was assigned to the pair of residues with the closest atom-atom pair and all other interactions were considered hydrogen bonds, ensuring only one salt bridge was recorded per positive/negative charge pair.

### Supplementary Molecular Dynamics Methods

**TCR:pMHC and TCRm:pMHC structure preparation.** X-ray crystal structures of four pMHC-TCRm and eight pMHC-TCR complexes were obtained from the Protein Data Bank (1). Missing loops and heavy atoms were rebuilt using CHARMM35 (2) based on SEQRES records and default residue parameters (see SI Table (SI Table 4)); short sections of missing residues at N- and C-termini were not rebuilt. Despite the increased simulation cost, TCR and TCR-mimetic antibody (TCRm) constant regions were included as this was recommended by previous studies (3). The H++ webserver (4) (http://biophysics.cs.vt.edu/H++) was used to determine protonation states for all titratable groups in these rebuilt structures assuming a pH of 7 and internal and external dielectric constants of 10 and 80, respectively, at a salt concentration of 0.15 M. Amber-format parameter/topology and coordinate files were prepared for periodic boundary simulations with tleap (5) using FF14SB (6) and TIP3P (7) forcefield parameters for protein and water, respectively. The PBRadii setting mbondi2 was used to prepare all solvated and dry topology files. Each complex was solvated in an orthorhombic box with a minimum distance between protein atoms and the box edge of 1.4 nm. Sodium and chloride ions were then added to neutralize each system and bring the salt concentration to 0.15 M, with ion counts chosen using the SPLIT method (8).

**Molecular dynamics simulation preparation.** All minimization, heating, and production simulations were carried out using OpenMM v7.5 (9). In all simulations, bonds containing hydrogen were constrained, the particle-mesh Ewald method (10) was used to compute long-range electrostatic interactions, and a non-bonded cutoff of 1 nm was used to compute short-range non-bonded interactions. All NVT simulations were conducted with a Langevin Middle Integrator (11) with a collision frequency of 1 $ps^{-1}$ and integration time step of 2 fs. NPT simulations were conducted identically to NVT simulations but with the addition of a Monte Carlo barostat (updated every 10 integration time steps) to maintain system pressure at 1 bar. The rebuilt, solvated, and neutralized structures were first minimized for 1,000 iterations with spherical harmonic restraints on all protein heavy atoms with force constants of 10 kcal/[mol x $Å^2$]. Systems were then heated from 48 to 298 K in steps of 10 K in the NVT ensemble for 10 ps at each temperature with all protein heavy atom restraints maintained. After heating, another round of 1,000 iterations of minimization was performed, with spherical harmonic restraints with force constant 5 kcal/[mol x $Å^2$] on all protein $C_\alpha$ atoms. A second round of heating was then carried out, again in the NVT ensemble, from 48 to 298 K in steps of 10 K with $C_\alpha$ atom restraints maintained and 2.5 ps of dynamics simulated at each temperature. Spherical harmonic restraints were then sequentially relaxed and the system density allowed to equilibrate during a series of 100-ps NPT simulations at 298 K with $C_\alpha$ atom restraint force constants of 5, 4, 3, 2, 1, 0 kcal/[mol x Å$^2$] with the final 100 ps of equilibration unrestrained.

**Production simulations, MMGBSA calculations, and contact analysis.** Production trajectories of 5-ns duration in the NPT ensemble were initiated from the final coordinates of the unrestrained equilibration. A total of thirty statistically independent trajectories were each run through this minimization, heating, equilibration, and simulation procedure for each pMHC-TCR or pMHC-TCRm system. A total of 3,000 simulation frames collected at 40-ps intervals within the final 4 ns of each 5-ns production trajectory (i.e., 100 frames from each of 30 trajectories per complex) were analyzed using the MMPBSA.py (12) program from AmberTools21 (5). Complex, receptor, and ligand trajectories were all collected from simulations of the bound complex conducted as described above. Generalized-Born implicit solvent (13) calculations were carried out with igb 2 (compatible with mbondi2) and saltcon set to 0.15. In all calculations the receptor was defined as the MHC Class I and beta-2-microglobulin chains plus the peptide antigen, while the ligand was defined as the two TCR or TCRm chains. MMGBSA energies were decomposed over all residues using the MMPBSA.py DECOMP functionality with idecomp set to 1. The fraction of the DECOMP energy, $f_{\mathrm{DECOMP}}$, contributed by the peptide, MHC alpha 1 ($\alpha1$) helix, or MHC alpha 2 ($\alpha2$) helix was computed as:

$$f_{DECOMP}^{pep} = \frac{\sum^{pep} g(i)}{\sum^{pep} g(i) + \sum^{\alpha1} g(i) + \sum^{\alpha2} g(i)}, \quad (1)$$

$$f_{DECOMP}^{\alpha1} = \frac{\sum^{\alpha1} g(i)}{\sum^{pep} g(i) + \sum^{\alpha1} g(i) + \sum^{\alpha2} g(i)}, \quad (2)$$

$$f_{DECOMP}^{\alpha2} = \frac{\sum^{\alpha2} g(i)}{\sum^{pep} g(i) + \sum^{\alpha1} g(i) + \sum^{\alpha2} g(i)}, \quad (3)$$

respectively. In these equations, $g(i)$ is the MMGBSA free energy change for residue $i$, and $\sum^{pep} g(i)$, $\sum^{\alpha1} g(i)$, and $\sum^{\alpha2} g(i)$ are summations over the per-residue free energy changes for all residues in the peptide antigen, $\alpha1$ helix, and $\alpha2$ helix, respectively. Fractional contributions from the CDRs of a TCR or TCRm were calculated in an analogous fashion, with the numerator a summation of the $g(i)$ values for a particular CDR and the denominator replaced by the sums of the energies over all six CDRs in the complex.

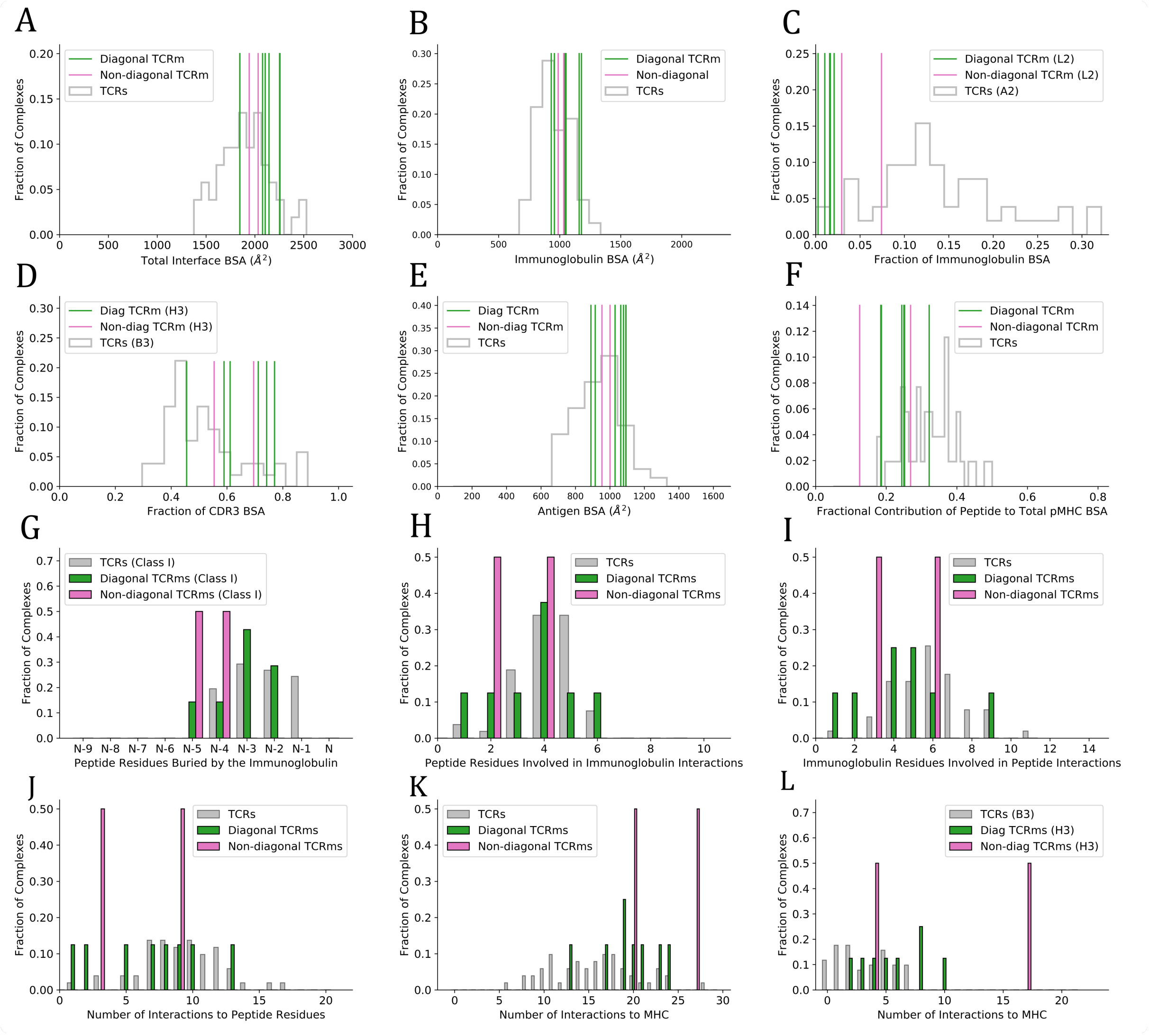


**Fig. 1.** pMHC engagement properties of diagonal (green lines/bars) and non-diagonal (pink lines/bars) TCRms, in the context of the properties of representative TCRs (grey histograms). A: The total BSA across the immunoglobulin:antigen interface. B: The immunoglobulin portion of the total interface BSA. C: The fractional contribution of CDRL2 (TCRms) or CDRA2 (TCRs) to immunoglobulin BSA. D: The fractional contribution of CDRH3 (TCRms) or CDRB3 (TCRs) to CDR3 BSA. E: The pMHC portion of the total interface BSA. F: The fractional contribution of peptide BSA to pMHC BSA. G: The number of peptide residues buried in each immunoglobulin:pMHC Class I complex, expressed in terms of the total length of the peptide, N. The peptides range from N = 8 to N = 10 residues in length. H: The number of peptide residues involved in binding interactions to the TCRms/TCRs. I: The number of immunoglobulin residues involved in binding interactions to the peptide across the TCRms/TCRs. J: The number of interactions between the immunoglobulin and the peptide across the TCRms/TCRs. K: The number of interactions to MHC residues across the TCRms/TCRs. L: The number of interactions to MHC residues made by the CDRH3 (TCRms) or CDRB3 (TCRs) loop.

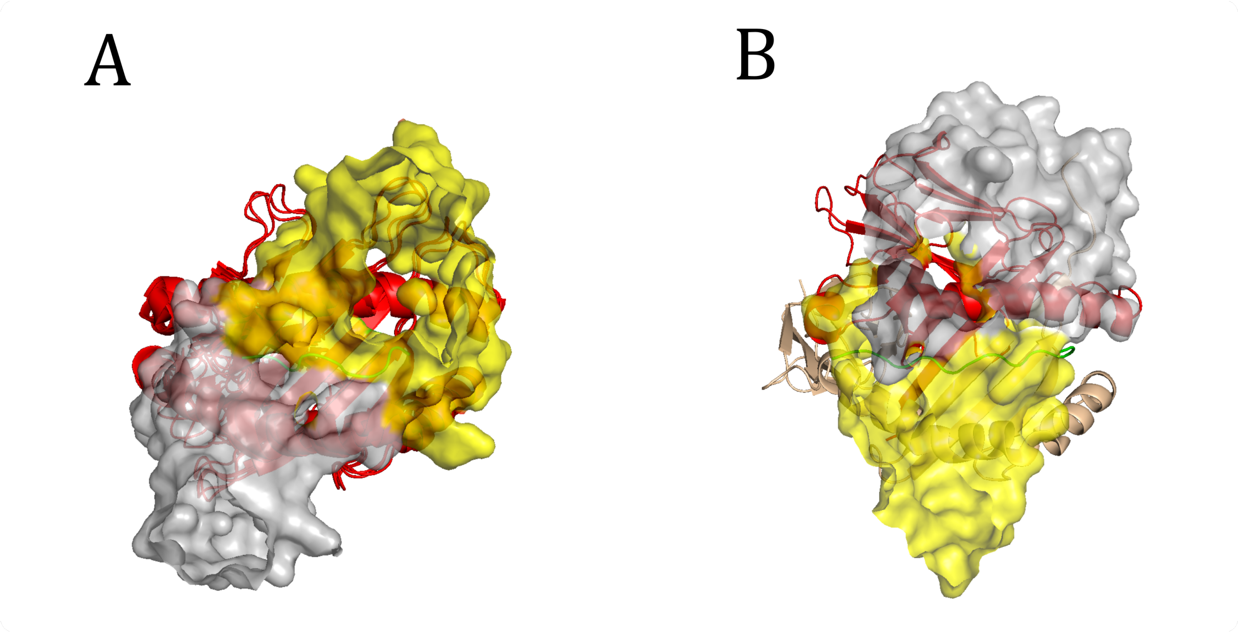


**Fig. 2.** TCR diagonal binding modes to peptide:MHC. A: the canonical diagonal TCR:pMHC binding mode (here illustrated by 5WLG in the context of a class I pMHC), B: the 'reverse-polarity' diagonal TCR:pMHC binding mode (here illustrated by 4Y19 in the context of a class II pMHC). Both exhibit similar absolute docking angles but with the roles of the TCR beta (VB) and alpha (VA) chains reversed. The non-canonical mode (B) is linked with weak/no T-cell signalling (14). Red cartoon: MHC alpha chain; wheat cartoon: MHC beta chain; green cartoon: peptide; yellow surface: IMGT-defined VB (15), light gray surface: IMGT-defined VA (15).

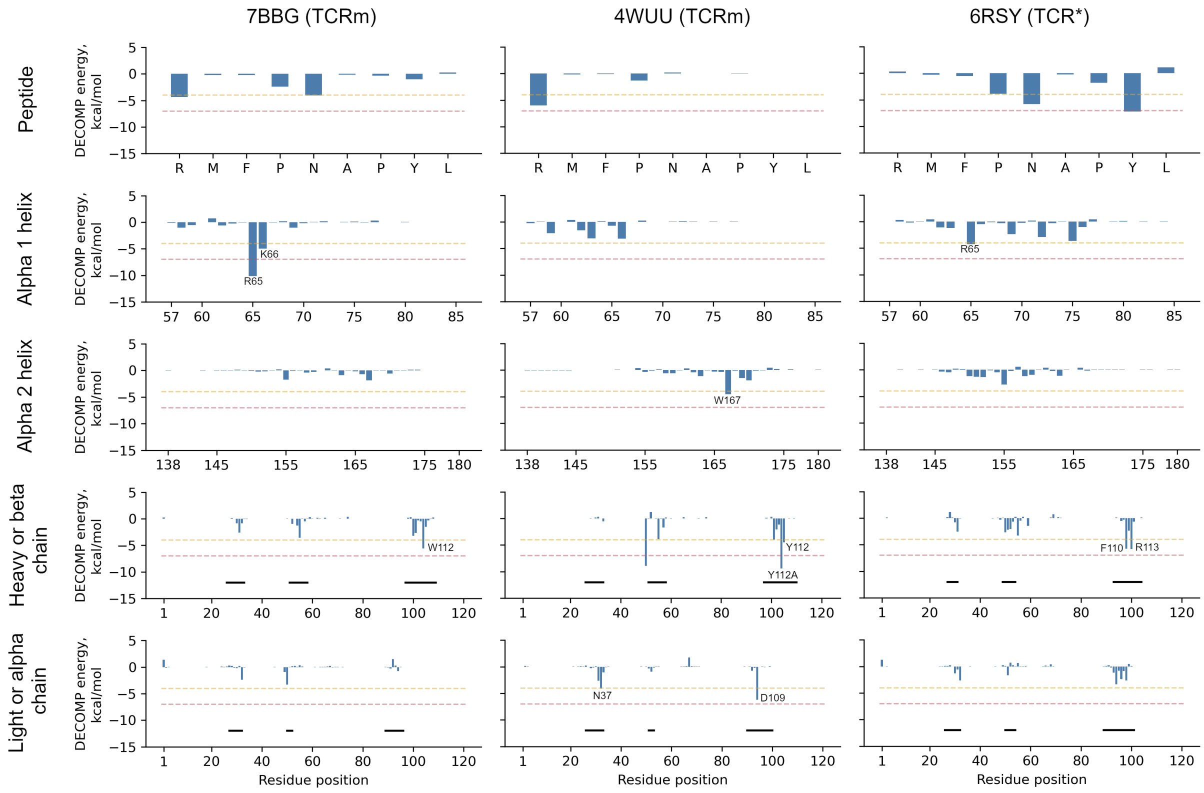


**Fig. 3.** MMGBSA decomposition results (12) for all Wilms Tumor 1 pMHC-TCRm/TCR complexes. For TCR or TCR mimetic chains, x-axis labels give residue positions numbered sequentially from 1 while labels on individual peaks refer to CDR loop numbers generated with ANARCI (16) using the IMGT (15) scheme. Horizontal black bars indicate the locations of, from left to right, CDR1, CDR2, and CDR3 within the sequence. The dashed amber and red lines reflect the energy thresholds to be considered 'semi-hotspots' (-4 kcal/mol) and 'hotspots' (-7 kcal/mol), respectively. *Affinity-enhanced TCR. PDB code to immunoglobulin name mappings — 4WUU: ESK1, 6RSY: a7b2, 7BBG: 11D06.

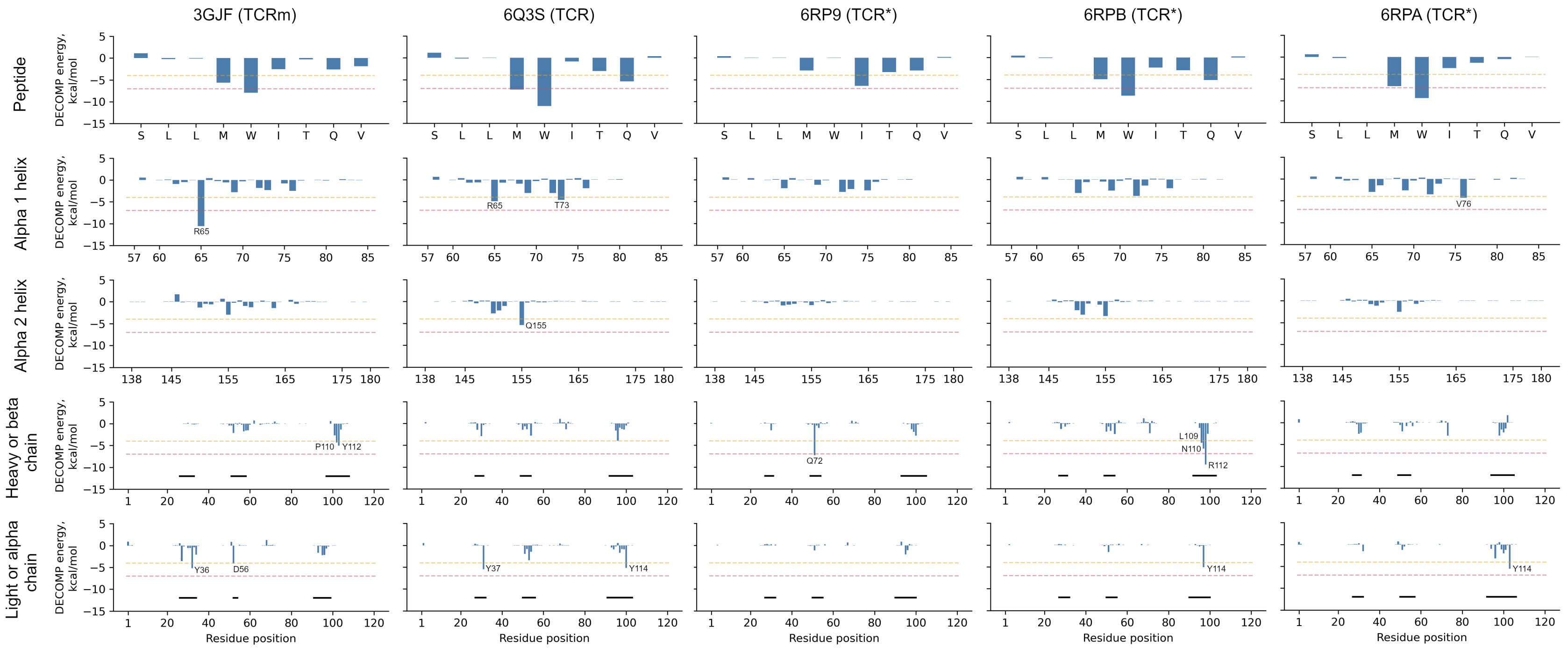


**Fig. 4.** MMGBSA decomposition results (12) for all New York esophageal squamous cell carcinoma 1 (NY-ESO-1) pMHC-TCRm/TCR complexes. For TCR or TCR mimetic chains, x-axis labels give residue positions numbered sequentially from 1 while labels on individual peaks refer to CDR loop numbers generated with ANARCI (16) using the IMGT scheme (15). Horizontal black bars indicate the locations of, from left to right, CDR1, CDR2, and CDR3 within the sequence. The dashed amber and red lines reflect the energy thresholds to be considered ‘semi-hotspots’ (-4 kcal/mol) and ‘hotspots’ (-7 kcal/mol), respectively. *Affinity-enhanced TCR. PDB code to immunoglobulin name mappings — 3GJF: 3M4E5, 6Q3S: sp3.4, 6RPA: NYE_S2, 6RPB: NYE_S1, 6RP9: NYE_S3.

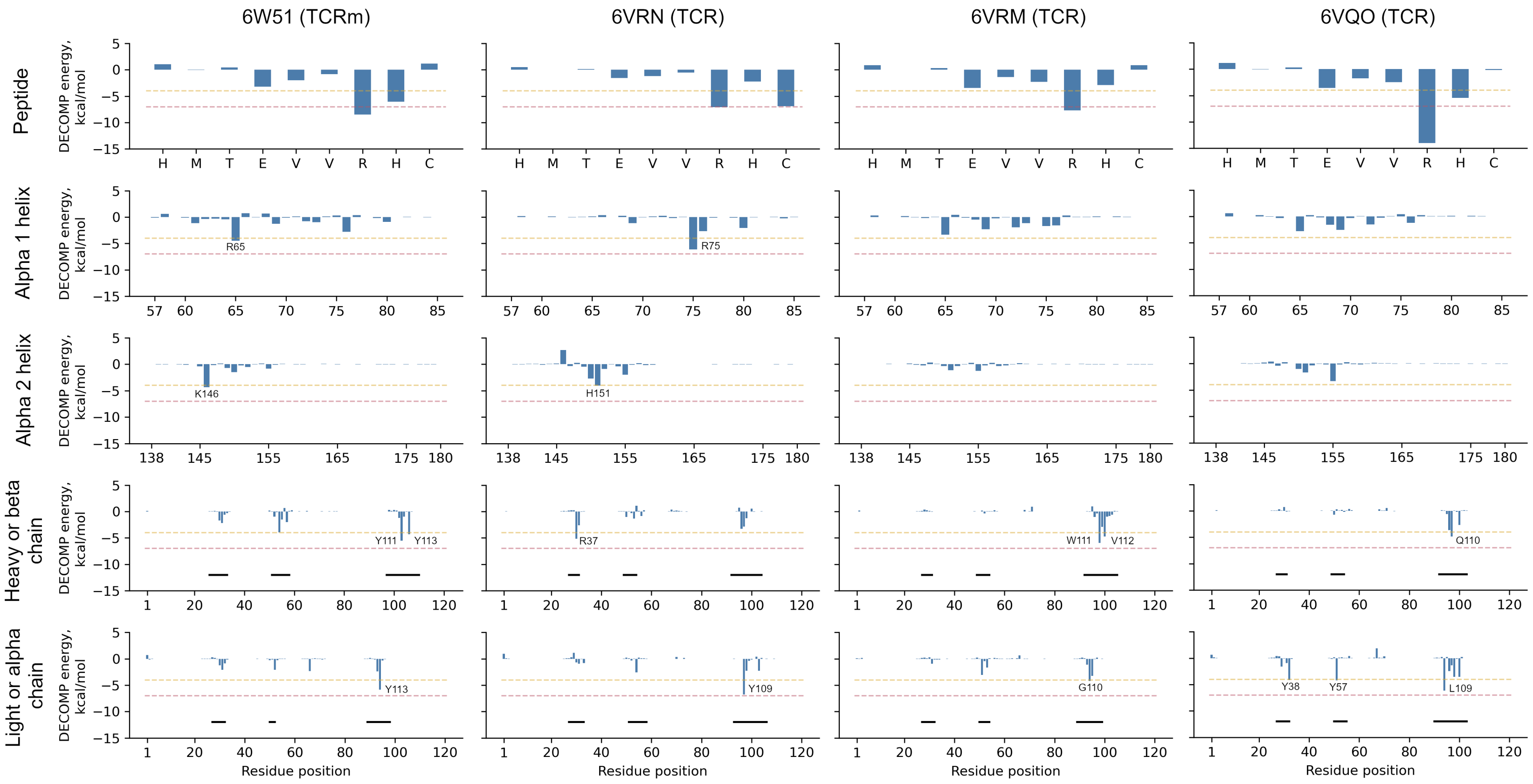


**Fig. 5.** MMGBSA decomposition results (12) for all p53 R175H neoantigen (p53_R175H) pMHC-TCRm/TCR complexes. For TCR or TCR mimetic chains, x-axis labels give residue positions numbered sequentially from 1 while labels on individual peaks refer to CDR loop numbers generated with ANARCI (16) using the IMGT scheme (15). Horizontal black bars indicate the locations of, from left to right, CDR1, CDR2, and CDR3 within the sequence. The dashed amber and red lines reflect the energy thresholds to be considered 'semi-hotspots' (-4 kcal/mol) and 'hotspots' (-7 kcal/mol), respectively. PDB code to immunoglobulin name mappings — 6VQO: 1a2, 6VRM: 12-6, 6VRN: 38-10, 6W51: H2.

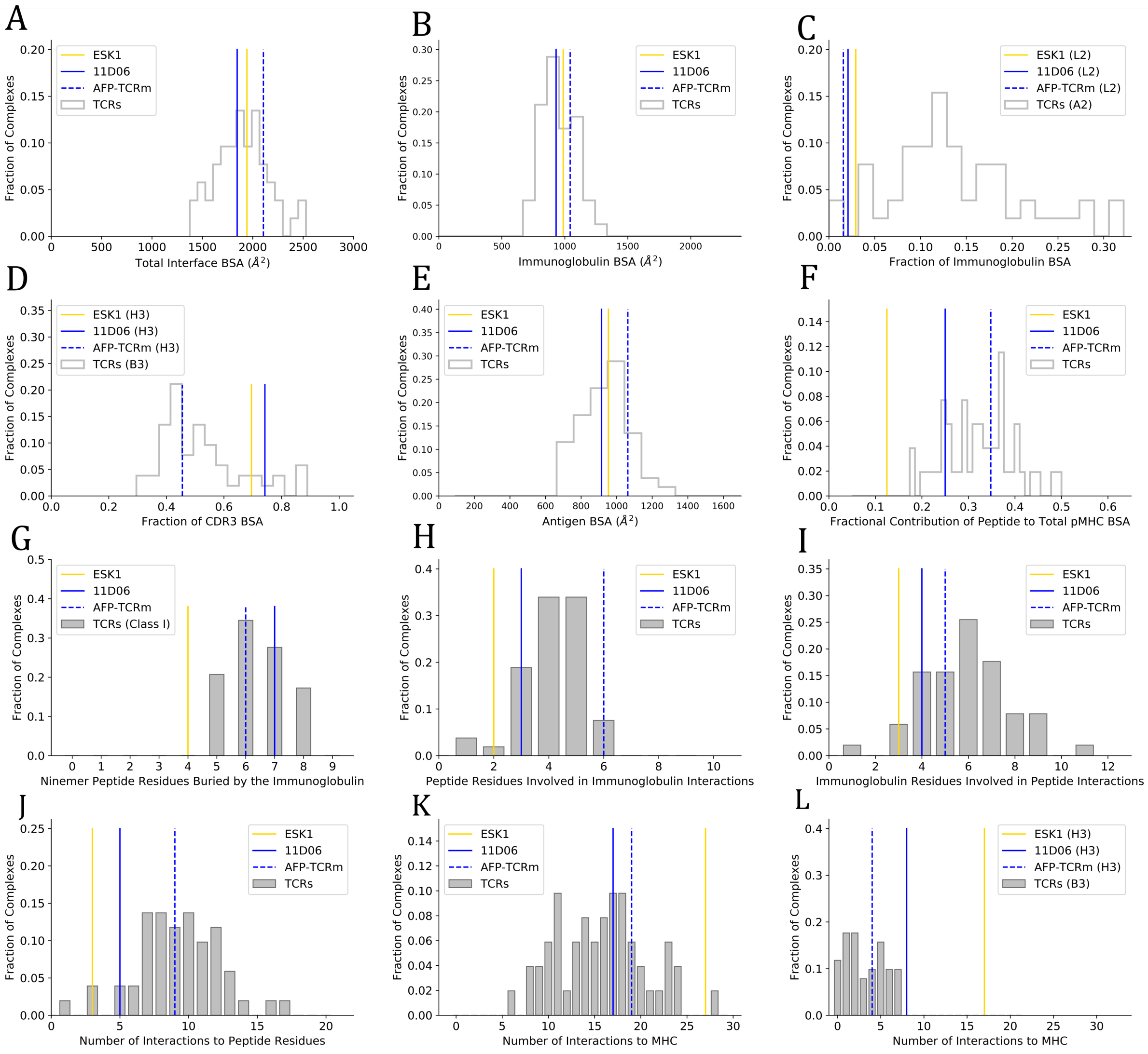


**Fig. 6.** pMHC engagement properties of two TCRms that are currently in clinical trials (11D06 [blue solid line] and AFP-TCRm [blue dashed line]) and a TCRm suspended at the preclinical stage (ESK1 [yellow solid line]), in the context of the properties of representative TCRs [grey histograms/bars]. ESK1 and 11D06 (*i.e.* both solid lines) engage the same pMHC complex (WT1 + HLA-A*02:01). A: The total BSA across the immunoglobulin:antigen interface. B: The immunoglobulin portion of the total interface BSA. C: The fractional contribution of CDRL2 (TCRms) or CDRA2 (TCRs) to immunoglobulin BSA. D: The fractional contribution of CDRH3 (TCRms) or CDRB3 (TCRs) to CDR3 BSA. E: The pMHC portion of the total interface BSA. F: The fractional contribution of peptide BSA to pMHC BSA. G: The number of peptide residues buried in each immunoglobulin to ninemer peptide:MHC Class I complexes. H: The number of peptide residues involved in binding interactions to the TCRms/TCRs. I: The number of immunoglobulin residues involved in binding interactions to the peptide across the TCRms/TCRs. J: The number of interactions between the immunoglobulin and the peptide across the TCRms/TCRs. K: The number of interactions to MHC residues across the TCRms/TCRs. L: The number of interactions to MHC residues made by the CDRH3 (TCRms) or CDRB3 (TCRs) loop.

| Interaction Type | Proportion in TCR:pMHC Interfaces ($\mu \pm$ sd) | Proportion in TCRm:pMHC Interfaces ($\mu \pm$ sd) |
|---|---|---|
| Hydrophobic | 56.3% ± 10.2% | 54.2% ± 8.5% |
| Aromatic | 4.0% ± 4.3% | 4.2% ± 3.9% |
| Polar | 39.7% ± 10.7% | 41.6% ± 8.0% |

**Table 1.** The proportions of each interaction type seen across the TCR:pMHC and TCRm:pMHC interfaces. Hydrogen bonds and salt bridges are pooled as 'polar' interactions.

| Name (PDB) | CDRH1 (Length) | CDRH2 (Length) | CDRH3 (Length) | CDRL1 (Length) | CDRL2 (Length) | CDRL3 (Length) |
|---|---|---|---|---|---|---|
| Hyb3 (1W72) | GFTFDDYA (8) | ISWNSGSI (8) | ARGRGFHYYYYGMDI (15) | NIGSRS (6) | DDS (3) | QVWDSRTDHWV (11) |
| 25-D1.16 (3CVH) | GYTFTDYN (8) | INPNNGGT (8) | ARKPYYGNFAWFAY (14) | EDIYNR (6) | GAT (3) | QQYWSTPLT (9) |
| 3M4E5 (3GJF) | GFTFSTYQ (8) | IVSSGGST (8) | AGELLPYYGMDV (12) | SRDVGGYNY (9) | DVI (3) | WSFAGSYYV (9) |
| 3M4F4 (3HAE) | GFTFSAYG (8) | IGSSGGGT (8) | AGELLPYYGMDV (12) | SRDVGGYNY (9) | DVI (3) | WSFAGSYYV (9) |
| ESK1 (4WUU) | GYSFTNFW (8) | VDPGYSYS (8) | ARVQYSGYYDWFDP (14) | SSNIGSNT (8) | SNN (3) | AAWDDSLNGWV (11) |
| 2Q1 (6UJ9) | GFNVKYYM (8) | ISPGYDYT (8) | SRSYWRYSVDV (11) | QDVNTA (6) | SAS (3) | QQVYSSPFT (9) |
| H2 (6W51) | GFNVYASG (8) | IYPDSDYT (8) | SRDSSFYYVYAMDY (14) | QDVNTA (6) | SAY (3) | QQYSRYSPVT (10) |
| 11D06 (7BBG) | GGTFSSYA (8) | IIPIFGTA (8) | ARSIELWWGGFDY (13) | QSISSW (6) | DAS (3) | QQYEDYTT (8) |
| 3H4 (7BH8) | GYTFTDYN (8) | INPNNGGT (8) | ARPDYYGSSYGWYFDV (16) | QDINSY (6) | RAN (3) | LQYDEFPLT (9) |
| AFP-TCRm* (7RE7) | GYSFPNYW (8) | IDPGDSYT (8) | ARYYVSLVDI (10) | SSDVGGYNY (9) | DVN (3) | SSYTTGSRAV (10) |
| 3.C11 (6XP6) | GGTVRSRVHA (10) | IIPIFGTA (8) | ARDVQRMGMDV (11) | QDISNW (6) | DSS (3) | QQFNSYPLT (9) |

**Table 2.** IMGT-defined (15) Complementarity-Determining Region (CDR) properties of the full set of TCR-mimetic antibodies (TCRms). All bind a peptide presented by Class-I MHC, except 3.C11 which recognises a peptide presented by Class-II MHC. *AFP-TCRm refers to the unnamed clinical-stage TCRm from Liu *et al.* 2022 (17).

| Antigen | Ig Name [PDB ID] | TCRm/TCR | Chain | CDRH1 Profile | CDRH2 Profile | CDRH3 Profile |
|---|---|---|---|---|---|---|
| WT1 | ESK1 [4WUU] | TCRm | VH | GYSFTNFW | VDPGYSYS | ARVQYSGYYDWFDP |
| | | | VL | SSNIGSNT | SNN | AAWDDSLNGWV |
| | 11D06 [7BBG] | TCRm | VH | GGTFSSYA | IIPIFGTA | ARSIELWWGGFDY |
| | | | VL | QSISSW | DAS | QQYEDYTT |
| | a7b2 [6RSY] | TCR* | VB | SEHNR | FQNEAQ | ASSLGFGRDVMR |
| | | | VA | TVDPNEY | GLKNN | IGGGTTSGTYKYI |
| NYESO-1 | 3M4E5 [3GJF] | TCRm | VH | GFTFSTYQ | IVSSGGST | AGELLPYYGMDV |
| | | | VL | SRDVGGYNY | DVI | WSFAGSYYV |
| | sp3.4 [6Q3S] | TCR | VB | MNHEY | SVGAGI | ASSYVGNTGELF |
| | | | VA | DSAIYN | IQSSQRE | AVRPTSGGSYIPT |
| | NYE_S3 [6RP9] | TCR* | VB | SGHVS | FNYEAQ | ASSSPGGVSTEAF |
| | | | VA | DRGSQS | IYSNGD | ALTRGPGNQFY |
| | NYE_S2 [6RPA] | TCR* | VB | SQVTM | ANQGSEA | SVGGSGGADTQY |
| | | | VA | VSGNPY | YITGDNLV | AVRDINSGAGSYQLT |
| | NYE_S1 [6RPB] | TCR* | VB | MNHEY | SVGAGI | ASSYLNRDSALD |
| | | | VA | DRGSQS | IYSDGD | AVKSGGSYIPT |
| p53_R175H | H2 [6W51] | TCRm | VH | GFNVYASG | IYPDSDYT | SRDSSFYYVYAMDY |
| | | | VL | QDVNTA | SAY | QQYSRYSPVT |
| | 12-6 [6VRM] | TCR | VB | MNHNS | SASEGT | ASSEGLWQVGDEQY |
| | | | VA | NSASQS | VYSSG | VVQPGGYQKVT |
| | 38-10 [6VRN] | TCR | VB | ENHRY | SYGVKD | AISELVTGDSPLH |
| | | | VA | TSENNYY | QEAYKQQN | AFMGYSGAGSYQL |
| | 1a2 [6VQO] | TCR | VB | MNHEY | SMNVEV | ASSIQQGADTQY |
| | | | VA | NSAFQY | TYSSGN | AMSGLKEDSSYKLI |

**Table 3.** Decomposed per-residue energetic profiles throughout the IMGT-defined (15) CDRs for the case study TCR and TCRm complexes. ‘Hotspots’ are defined as residues predicted to have an attractive per-residue contribution of ≥ 7 kcal/mol to free energy based on DECOMP analysis (underlined bold red text). ‘Semi-hotspots’ are defined as residues that contribute between -4 and -7 kcal/mol (underlined bold black text). *Affinity-enhanced TCRs. Ig: Immunoglobulin.

| PDB ID [Ig Name] | Chains Used | Residues Rebuilt (sequential numbering) |
|---|---|---|
| 3gjf [3M4E5] | D, E, F, K, M | None |
| 4wuu [ESK1] | A, B, C, D, E | None |
| 6q3s [sp3.4] | A, B, C, D, E | None |
| 6rp9 [NYE_S3] | A, B, C, D, E | None |
| 6rpa [NYE_S2] | A, B, C, D, E | Chain D (145-147, 160-166, 180-184), Chain E (236-240) |
| 6rpb [NYE_S1] | A, B, C, D, E | Chain A (221-225), Chain D (141-146) |
| 6rsy [a7b2] | A, B, C, D, E | None |
| 6vrm [12-6] | A, B, D, E, P | Chain D (1, 122, 125-131, 177-184) |
| 6vrn [38-10] | A, B, D, E, P | Chain D (1, 56-61) |
| 6vqo [1a2] | A, B, D, E, P | Chain D (1, 129-133, 182-184) |
| 6w51 [H2] | D, E, F, O, P | None |
| 7bbg [11D06] | A, B, C, H, L | Chain H (1, 135-141) |

**Table 4.** Chains used in the molecular simulations and residue regions rebuilt prior to simulation. Ig: Immunoglobulin.
